# Deep learning-based segmentation of high-resolution computed tomography image data outperforms commonly used automatic bone segmentation methods

**DOI:** 10.1101/2021.07.27.453890

**Authors:** Daniella M. Patton, Emilie N. Henning, Rob W. Goulet, Sean K. Carroll, Erin M.R. Bigelow, Benjamin Provencher, Nicolas Piché, Mike Marsh, Karl J. Jepsen, Todd L. Bredbenner

**Affiliations:** Department of Orthopaedic Research, University of Michigan, Ann Arbor, MI 48109; Mechanical and Aerospace Engineering Department, University of Colorado Colorado Springs, Colorado Springs, CO, 80918; Object Research Systems, Montreal, QC, CA

**Keywords:** Bone segmentation, threshold, deep learning, CNNs

## Abstract

Segmenting bone from background is required to quantify bone architecture in computed tomography (CT) image data. A deep learning approach using convolutional neural networks (CNN) is a promising alternative method for automatic segmentation. The study objectives were to evaluate the performance of CNNs in automatic segmentation of human vertebral body (micro-CT) and femoral neck (nano-CT) data and to investigate the performance of CNNs to segment data across scanners.

Scans of human L1 vertebral bodies (microCT [North Star Imaging], n=28, 53μm^3^) and femoral necks (nano-CT [GE], n=28, 27μm^3^) were used for evaluation. Six slices were selected for each scan and then manually segmented to create ground truth masks (Dragonfly 4.0, ORS). Two-dimensional U-Net CNNs were trained in Dragonfly 4.0 with images of the [FN] femoral necks only, [VB] vertebral bodies only, and [F+V] combined CT data. Global (i.e., Otsu and Yen) and local (i.e., Otsu r = 100) thresholding methods were applied to each dataset. Segmentation performance was evaluated using the Dice coefficient, a similarity metric of overlap. Kruskal-Wallis and Tukey-Kramer post-hoc tests were used to test for significant differences in the accuracy of segmentation methods.

The FN U-Net had significantly higher Dice coefficients (i.e., better performance) than the global (Otsu: p=0.001; Yen: p=0.001) and local (Otsu [r=100]: p=0.001) thresholding methods and the VB U-Net (p=0.001) but there was no significant difference in model performance compared to the FN + VB U-net (p=0.783) on femoral neck image data. The VB U-net had significantly higher Dice coefficients than the global and local Otsu (p=0.001 for both) and FN U-Net (p=0.001) but not compared to the Yen (p=0.462) threshold or FN + VB U-net (p=0.783) on vertebral body image data.

The results demonstrate that the U-net architecture outperforms common thresholding methods. Further, a network trained with bone data from a different system (i.e., different image acquisition parameters and voxel size) and a different anatomical site can perform well on unseen data. Finally, a network trained with combined datasets performed well on both datasets, indicating that a network can feasibly be trained with multiple datasets and perform well on varied image data.

## Introduction

Micro-computed tomography (micro-CT) imaging has become the “gold standard” for assessing bone morphology and microstructure in orthopedic research^1,2^. The first step in quantifying bone microstructure from micro-CT images involves separating bone voxels from background (i.e., non-bone) voxels and, typically, researchers use a thresholding approach^2–4^. However, descriptors of bone composition and architecture are sensitive to variations in the threshold values determined to separate bone from background^5–8^. For example, a 7.1% variation in a grey-level threshold can result in up to a 35 μm difference in trabecular thickness, which may directly affect study reproducibility and the biological interpretation of the data ^8^. Despite threshold sensitivity and the potential impact on study results, there is no standard method to segment bone from non-bone voxels ^2,7,8^.

For many imaging applications (e.g., fracture healing, growth and development), manual segmentation involves large labor costs to achieve accurate results ^2,7^. As a result, a number of global and local threshold methods have been used in bone-background segmentation. A global threshold involves choosing a single grayscale intensity value, typically from a bi-modal histogram ^9,10^. However, bone grayscale values may vary within and between scans because of mineral heterogeneity in the bone tissue or because of image noise, beam hardening, and scan-to-scan variability. These sources of grayscale variation limit the segmentation accuracy for global thresholding methods^2^. Alternatively, local thresholding (e.g., region growing and edge detection) determine a threshold for each voxel by considering neighboring voxels within a specified radius and allow the threshold value to change dynamically across the image ^11^. Local threshold methods often require setting a number of image-specific parameters, which may provide additional challenges ^11,12^. Consequently, bone research would benefit from the development of improved automated segmentation methods that can be broadly applied to bone scans with multiple sources of grayscale variation.

Recently, there has been rapid development of deep learning mathematical models such as convolutional neural networks (CNNs), which are feed-forward artificial neural networks developed specifically to tackle problems with image processing^13,14^. CNNs have been successfully used for classification, object detection, and segmentation of medical image data and have been shown to consistently outperform traditional approaches ^14^. In the musculoskeletal field, promising results have been shown using CNNs to segment vertebrae, whole body, and the proximal femurs in both clinical MR and CT scans ^15–19^. However, previous studies have created CNNs that perform well on one dataset^15–19^. In order for a network to be widely utilized for segmentation, a CNN must accurately and consistently segment bone tissue across a variety of micro-CT imaging data. While this has yet to be tested in segmentation tasks in bone research, a recent publication determined that a CNN can be trained to accurately classify medial temporal atrophy (i.e., a measure frequently used to measure dementia) from clinical magnetic resonance imaging data with variations in image quality^20^. Therefore, we hypothesized that a CNN could be trained and optimized to segment bone from a variety of micro-CT imaging data, thereby offering a more standardized approach to segment bone from background.

The first objective of this study was to create fully-connected CNNs (FCNNs) with U-net architecture to threshold bone from background in two separate datasets: [1] nano-CT scans of the femoral neck (27 μm isotropic voxels) and [2] micro-CT scans of the lumbar (L1) vertebral body (53 μm isotropic voxels). The U-Net architecture is particularly well adapted to perform well with small datasets, allowing network training with a limited number of manually segmented ground truth masks^21^. The resulting FCNNs were used to segment image data and the resulting masks were compared to those segmented using global or local thresholding methods. Finally, we investigated the performance of bone segmentation using a CNN trained using combined datasets [3]. We further hypothesized that the FCNN-based segmentation approach will significantly outperform threshold-based segmentation methods. The overarching goal of this study was to create an accurate segmentation method that performs well despite biological and system variability, resulting in greatly reduced time and effort required for segmentation and improved rigor and reproducibility.

## Methods

### Bone Samples

Fresh frozen cadaveric bone specimens were collected from adults with no observable or known musculoskeletal trauma or pathologies. Twenty-eight right proximal femurs [15 male, 60 ± 20 (18 - 89) years; 13 female, 61 ± 22 (24 - 95) years] were obtained from ScienceCare (Phoenix, AZ, USA), Anatomy Gifts Registry (Hanover, MD, USA), University of Michigan Anatomical Donations Program (Ann Arbor, MI, USA), and Ohio Valley Tissue and Skin Center (Cincinnati, OH, USA). Twenty-eight first lumbar (L1) vertebral bodies [14 male, 62.0 ± 15.2 (25 - 88) years; 14 female, 58.5 ± 14.3 (28 - 80) years] were obtained from The National Disease Research Interchange (Philadelphia, PA, USA). All samples were de-identified and considered unregulated human tissues (exemption 4) by the Institutional Review Boards.

### Sample Preparation and Scanning Protocol

Femurs were cut 16.5 cm below the superior aspect of the femoral head. Proximal femurs were positioned upright and scanned using a nano-computed tomography (nano-CT) system (phoenix nanotom-s, GE Measurement & Control; Wunstorf, Germany) and the following image acquisition parameters: 110 kV, 200 μA, 0.07 mm thick brass filter, 1 skips, 3 averages, 1000 images/360 degrees. Nano-CT image volumes were reconstructed with 27 μm dimensionally-isotropic voxels (phoenix datos|x, GE Sensing and Inspection Technologies; Wunstorf, Germany). Posterior processes were removed from the L1 vertebrae and the vertebral bodies were scanned using a micro-CT system (X50-CT, North Star Imaging, Rogers, MN, USA) with the following image acquisition parameters: 110 kV, 300 μA, continuous scan, 1080 images/360 degrees. Micro-CT image volumes were reconstructed with 53 μm dimensionally isotropic voxels (COBRA, Exxim Computing Corp.; Pleasonton, CA, USA).

### Image Processing and Generation of Ground Truth Data

Proximal femur images were converted to 16-bit intensity depth, image noise was removed using a 3D median filter (radius = 3) (MATLAB 2018a, The MathWorks Inc.; Natick, MA, USA), and femoral neck volumes were extracted (*JBMR manuscript in review*). Full vertebral body images were considered. Six transverse slices evenly spaced throughout the bone regions were selected in each dataset, resulting in a set of 168 nanoCT slices for the femoral neck and 168 microCT slices for the vertebral body. In the case of vertebral bodies, two of the six slices passed through the endplates. Images were cropped to contain only the bone data of interest. Otsu segmentation was used to generate initial bone masks from the image data. Bone masks were manually corrected with consensus of, at least, two (of three) individuals for bone voxels to generate ground truth segmentation masks (Dragonfly 4.0, Object Research Systems; Montreal, QC, Canada).

### Automatic Bone Segmentation Using a Deep Learning-Based (Supervised) Method

Intensity histograms for each image dataset were mapped to span the positive 16-bit integer range (0 – 32,767) to increase contrast and to account for variation in attenuation (grayscale levels) between bones and differences in the range of grayscale levels between imaging systems and between scans. Two-dimensional FCNNs were trained separately for the femoral neck and the vertebral body using a nested four-fold cross-validation approach (Fig. 1 a,b). Extracted image slices were divided by dataset into four folds of seven femoral necks and four folds of seven vertebral bodies, each containing 42 slices. Three folds (126 slices) of each bone type were combined for use as the training set. Thirteen slices from the training data (10%) were randomly selected as a validation set in Dragonfly (Dragonfly 4.0, Object Research Systems; Montreal, QC, Canada). A nested four-fold cross-validation approach was also used to train the FCNNs with the combined dataset (Fig. 1 c). Folds were combined for the vertebral body and femoral neck which resulted in each fold containing 84 slices, with each fold consisting of seven femoral necks and seven vertebral bodies.

**Figure 1:**
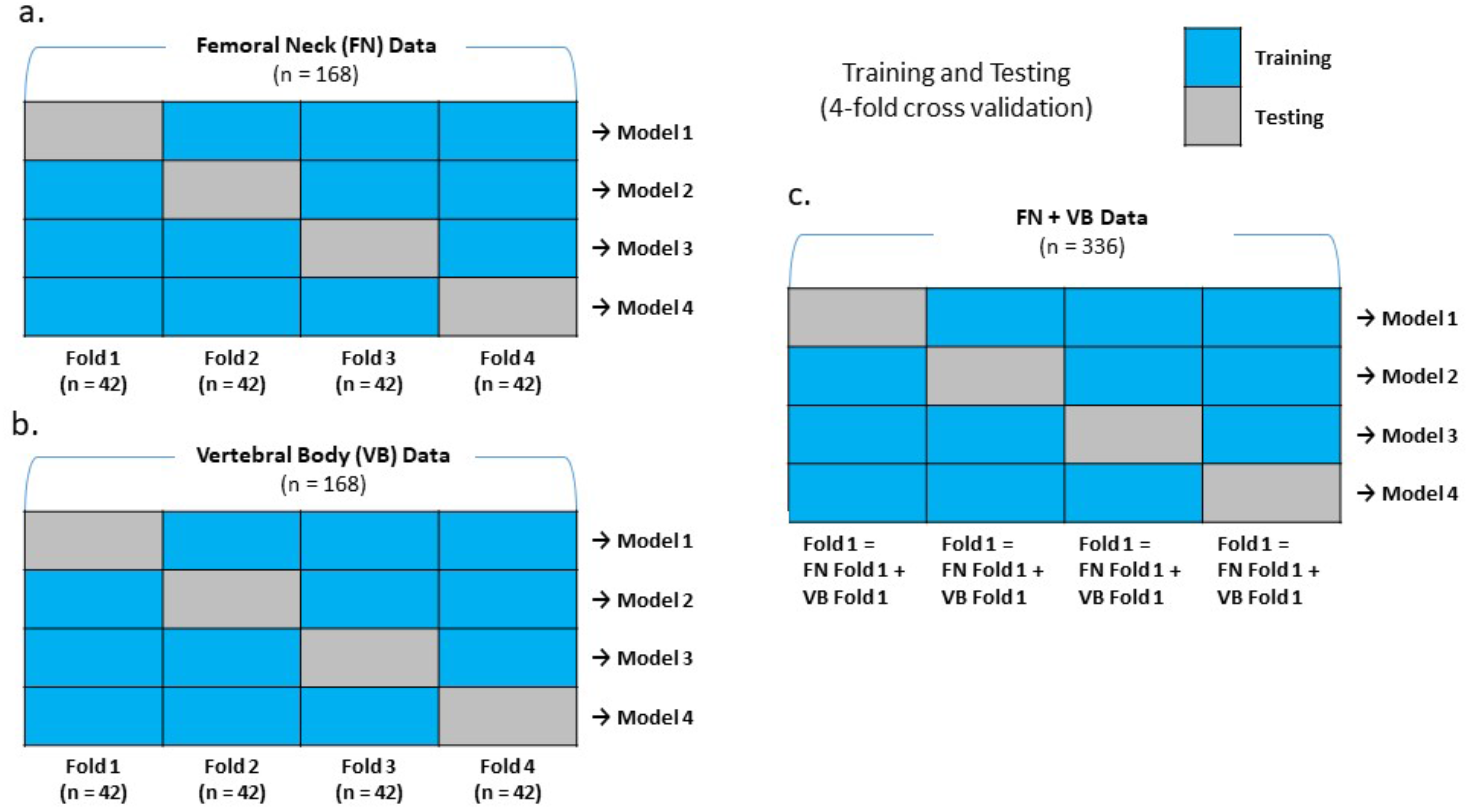
Schematic overview of the training and testing split using a nested 4-fold cross validation of the (a) vertebral body, (b) femoral neck, and (c) combined data FCNNs considered in this study.

Baseline 2D U-Net networks with the original U-Net weights and training parameters were optimized for the vertebral body and femoral neck datasets in this study ^21^. Network weights were determined using training set grayscale image data and ground truth masks with a batch size of 16, stride ratio of 1, patch size of 64, cross-entropy loss function, and Adadelta optimization (Dragonfly 4.0, Object Research Systems; Montreal, QC, CA). During training, the learning rate was adjusted as the optimizer reached a plateau by decreasing the step size by 10% with a minimum step size change of 0.1% and a patience parameter of 10. In addition, each model was trained for a maximum of 100 epochs or until the loss function did not decrease by more than 0.01% for 10 consecutive epochs. Test sets of the remaining fold of 42 slices from seven femoral necks or vertebral bodies were segmented using the resulting vertebral body FCNN (VB UNet) and femoral neck FCNN (FN UNet) on a slice-by-slice basis. This process was repeated four times, so each fold was used as a test set, resulting in four networks trained on the femoral neck and vertebral body data. Similarly, the FCNN trained with the combined vertebral body and femoral neck data (FN+VB UNet) was used to segment test sets (the combined fold of 84 slices from seven femoral necks and seven vertebral bodies) on a slice-by-slice basis. Accordingly, all image slices were segmented with a vertebral body FCNN, femoral neck FCNN, and a combined FCNN trained using.

The FCNNs trained with the vertebral body data were also tested on the femoral neck images, and the FCNNs trained with the femoral neck data were tested with the vertebral body images. To accomplish this, the FCNNs trained with the three labeled folds in one dataset was only tested on the labeled fold that was not used on the unlike data. For example, a femoral neck FCNN trained with folds 1, 2, and 3 was tested only on fold 4 of the vertebral body data-set and vice-versa. This ensured that results from unlike data can be compared directly to all other results in this study.

### Automatic Bone Segmentation Using Unsupervised Methods

Local and global threshold methods were selected from available methods in the *scikit-image* package written in Python^23^. All threshold methods considered were applied to the 16-bit femoral neck and vertebral body data on a slice-by-slice basis in Python (v3.5) using Anaconda^24^. While many threshold methods were compared (Supp. 1), only a few will be considered for the remainder of analyses. Two global threshold methods, the Otsu minimum two-class variance^25^ and Yen’s threshold method^26^, were compared to the U-net segmentation. The global threshold methods were selected because the Otsu method is frequently used in orthopedic research^2,27^ and because the Yen threshold method qualitatively outperformed all other threshold methods considered. A local Otsu method was also applied to all image data on a slice-by-slice basis using local neighborhood analyses with a radius of 100 voxels. The scripts used to create all threshold masks and compare the segmentation quality are available on Github (https://github.com/daniella-patton/Threshold_Unet_Comparisons).

### Statistical Analyses

All corresponding segmented and ground truth bone masks were compared using the Dice similarity index (Dice) to quantify the performance of medical imaging data segmentation ^28^. Dice values measure the overlap of the segmented mask with the ground truth mask and range from 0.0 (no overlap) to 1.0 (perfect overlap). A non-parametric Kruskal-Wallis test^29^ followed by a Nemenyi post-hoc test^30^ was used to test for significant differences between the pooled measures of FCNN segmentation performance and all other methods. A significance level of 5% was used for all comparisons^31^.

## Results

### Model 1: Segmentation of Femoral Neck Data

The Otsu and Yen global threshold methods had a Dice indexof 0.445 ± 0.130, and 0.894 ± 0.056 relative to the ground truth femoral neck data, respectively. The mean Dice index of 0.283 ± 0.089 for the local Otsu threshold was lower than either of the global threshold methods considered here. All U-Net models had average Dice indices greater than 0.89, indicating high similarity in segmentation quality relative to the ground truth data. The FN UNet had a Dice index of 0.966 ± 0.018, whereas the VB UNet had a DICE coefficient of 0.892 ± 0.080. Finally, the FN+VB UNet had a DICE coefficient of 0.963 ± 0.018. A boxplot and summary statistics are presented below (Fig. 2).

**Figure 2:**
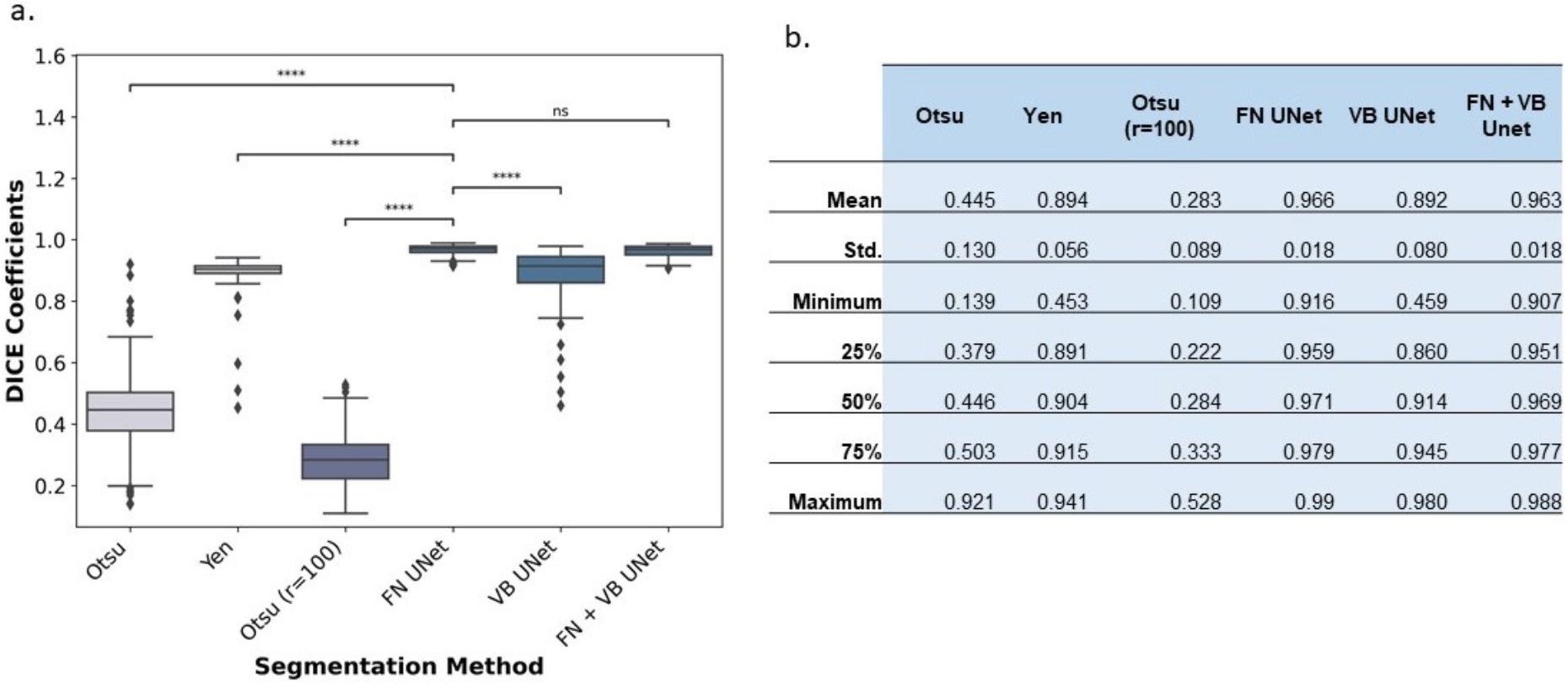
(a) A boxplot with a Nemenyi post-hoc comparison of the DICE coefficients calculated on a slice-by slice basis for the global, local, and previously trained U-net segmentation methods relative to the ground truth femoral neck dataset. Diamonds in the boxplot represent outliers in the data (**** p <= 0.001, ns p > 0.05). (b) A table with more specific statistics of the DICE coefficients of the femoral neck dataset (i.e., the mean, standard deviation [std.], minimum, 25%, 50%, 75%, and maximum) are summarized.

The Kruskal-Wallis test demonstrated that there are significant differences in the segmentation methods compared in this study (p <0.001). The Nemenyi post-hoc test revealed that the FN UNet had a significantly higher Dice index compared to the Yen (p=0.001), global Ostu (p=0.001), local Ostu (p=0.001), and VB UNet (p=0.001) (Figure 2). Finally, there was no significant difference between the Dice indices for FN UNet and the FN+VB UNet (p=0.783), indicating that including “un-like” data with training did not negatively affect model performance. Typical performance of each segmentation method relative to the ground truth data is presented below (Fig. 3).

**Figure 3:**
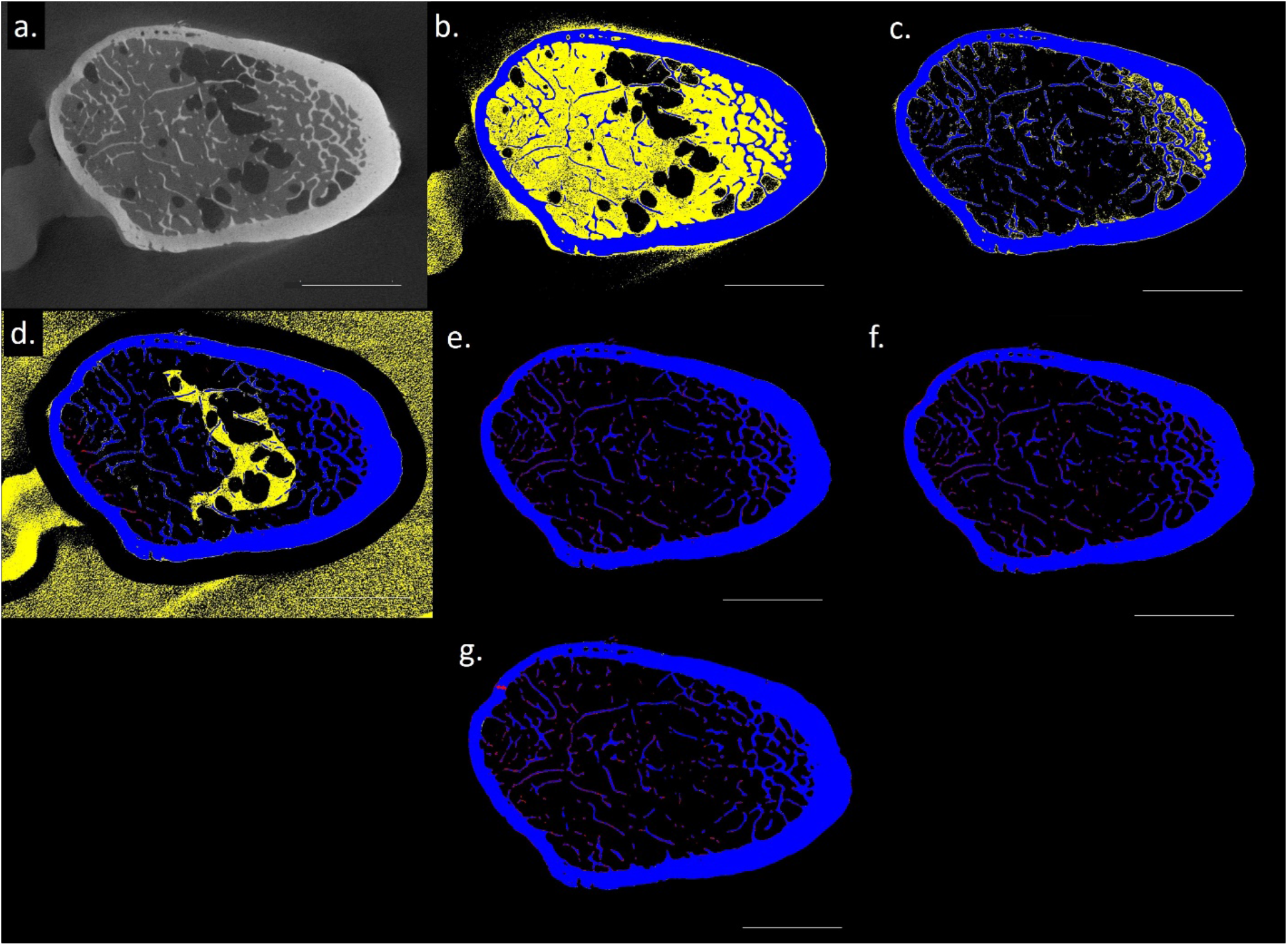
A visual representation of a femoral neck (a) input slice, (b) global Otsu threshold, (c) global Yen threshold, (d) local Otsu threshold (r=100), (e) FN UNet, (f) VB UNet, and (g) FN+VB UNet. For each mask presented the colors demonstrate where each method overestimates bone (yellow), underestimates bone (red), and correctly identifies bone (blue) in an individual femoral neck cross section (scale bar = 10 mm).

### Model 2: Segmentation of Vertebral Body Data

The Otsu and Yen global threshold methods had a Dice index of 0.356 ± 0.091 and 0.812 ± 0.220 relative to the ground truth vertebral body data, respectively. The mean Dice index of 0.444 ± 0.110 for the local Otsu threshold method is higher than the Dice index for the global Otsu threshold (0.356 ± 0.091) but lower than the Yen threshold (0.812 ± 0.220). The VB UNet had a Dice index of 0.881 ± 0.102 and the FN UNet had a Dice index of 0.777 ± 0.104. Finally, the FN+VB UNet had a Dice index of 0.912 ± 0.054. A boxplot and specific summary statistics are visually presented below (Figure 4).

**Figure 4:**
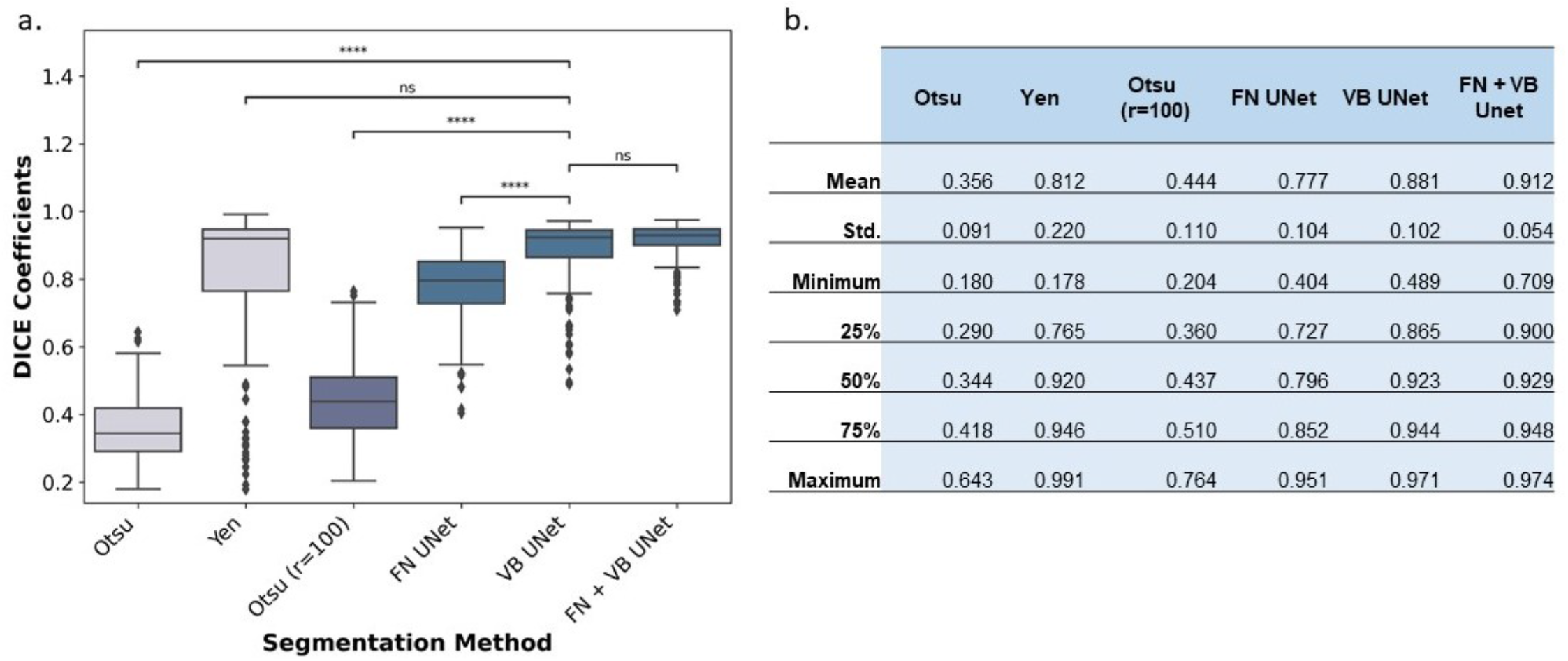
(a) A boxplot with a Nemenyi post-hoc comparison of the DICE coefficients calculated on a slice-by slice basis for the global, local, and previously trained U-net segmentation methods relative to the ground truth vertebral body datasets. Diamonds in the boxplot represent outliers in the data. (**** p <= 0.001, ns p > 0.05) (b) A table with more specific statistics of the DICE coefficients of the femoral neck dataset (i.e., the mean, standard deviation [std.], minimum, 25%, 50%, 75%, and maximum) are summarized.

The Kruskal-Wallis test demonstrated that there are significant differences in segmentation quality for the vertebral body dataset (p <0.001). The Nemenyi post-hoc test revealed that the VB UNet had a significantly higher Dice index compared to the global and local Otsu (p=0001 for both), but not compared to the Yen threshold (p=0.462) (Fig. 4). The VB UNet also had higher Dice indices compared to the FN UNet (p=0.001) but was not significantly different from the FN+VB UNet (p=0.616). As with training on femoral neck images, including un-like data with training did not significantly affect model performance. A visual example performance of each segmentation method relative to the ground truth data is presented below (Fig. 5).

**Figure 5:**
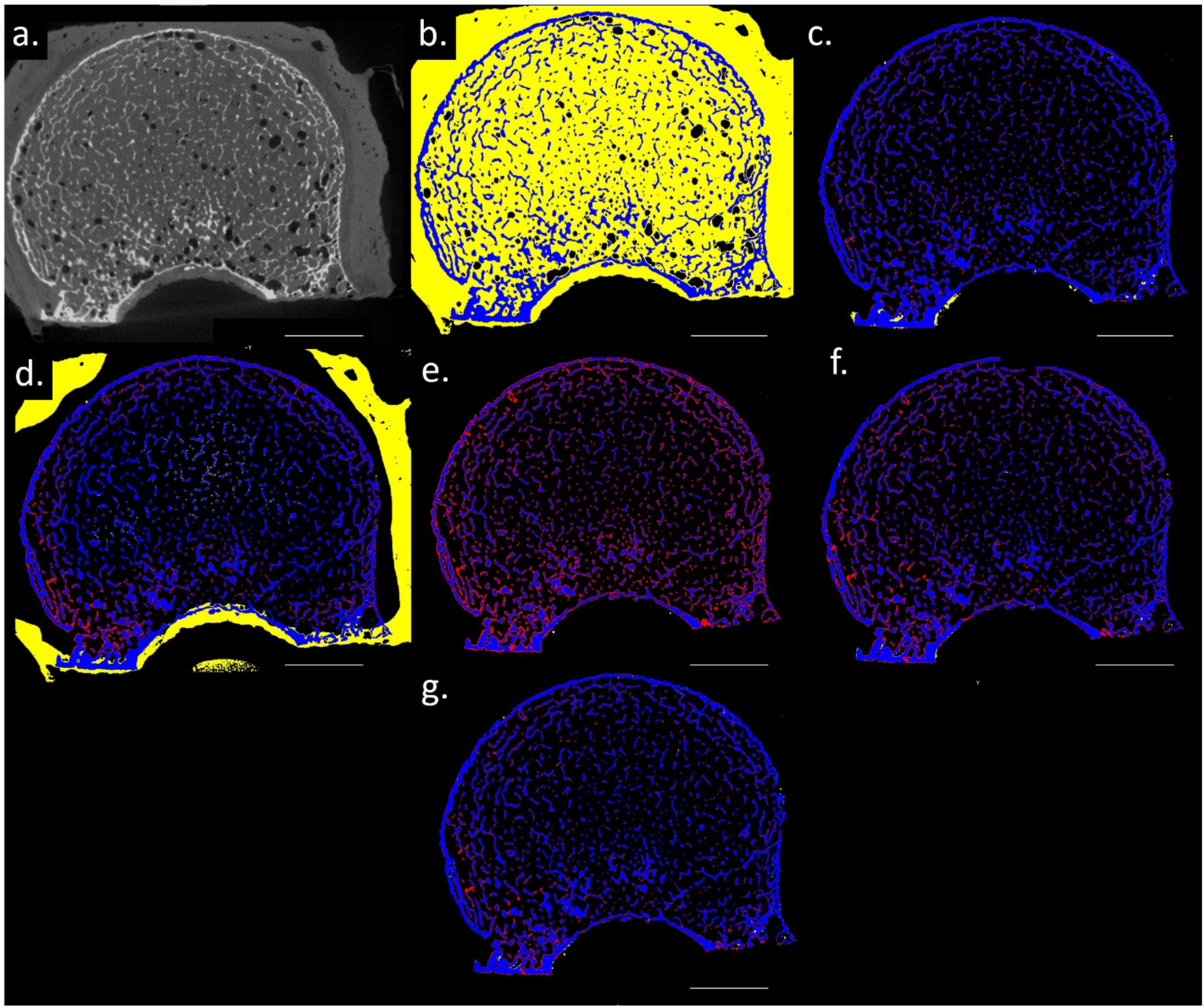
A visual representation of a vertebral body (a) input slice, (b) global Otsu threshold, (c) global Yen threshold, (d) local Otsu threshold (r=100), (e) FN UNet, (f) VB UNet, and (g) FN+VB UNet. For each mask presented the colors demonstrate where each method overestimates bone (yellow), underestimates bone (red), and correctly identifies bone (blue) in an individual vertebral body cross section (scale bar = 10 mm).

## Discussion

Segmenting bone from background in high resolution CT scans is a critical step in quantifying bone microarchitecture. Although automatic segmentation methods are used frequently, there is no single method that performs well when broadly applied across orthopedic research^2^. Our study confirmed that an FCNN can be trained to segment bone scans from imaging modalities frequently used in research at a higher level of performance than commonly-used thresholding methods. More importantly, these findings show that a single FCNN can be trained to perform well on images of different bones acquired from different image-acquisition systems and protocols. Currently, this combined network offers an immediate and alternative method for bone segmentation in CT scans (publicly available on the infinite toolbox in Dragonfly 4.0). These findings demonstrate that a trained network can perform well on datasets which vary in bone structure and acquisition parameters. The time involved in training the U-Nets was minimal and was justified given the significant improvement in performance. Once established, a U-Net is expected to reduce time and effort required for segmentation, while also improving rigor and reproducibility.

The FCNN’s trained on the vertebral body or femoral neck sections outperformed nearly all threshold methods considered in this study. The high performance of the FCNNs was achieved despite being considered “small” datasets in deep learning research^14^. For example, one of the most popular networks used for image classification, the ImageNet, was trained on 15 million labeled high-resolution images for classification^32^. From the perspective of basic and translational research, this is an unreasonable and unattainable dataset size for any study. However, our findings demonstrated that training a network is possible with a small number of image slices (i.e., 113 slices for the vertebral body or femoral neck only FCNNS) and results in improved accuracy. The U-Net, the 2015 ISBI cell tracking winner, has continually proven to yield more precise segmentations with very few training images due to its unique contracting and expanding structure that are combined so that a convolution layer can learn more precise outputs ^21^. As a result, these findings are in line with the large volume of recent studies demonstrating the utility of the U-Net for end-to-end segmentation in medical imaging research (i.e., cited 17,657 times, *google scholar*).

In the musculoskeletal field, FCNNs are also promising segmentation tools, performing well on clinical magnetic resonance (MR), CT, and X-ray scans for the proximal femur, cartilage and bone, and knee bone tumor identification, respectively^15,16,18,33^. Despite research on clinical applications, the current study is the first to demonstrate FCNN applicability in high-resolution ex-vivo CT scans, which are commonly used to assess disease progression and/or drug treatment responses in bone^2^. The results argue for expanded use of a CNN such as the U-Net in basic research due to the small datasets needed for training and would ensure the results presented are accurate, and from a segmentation standpoint, reproducible.

Bone segmentation often relies on a global threshold approach ^2,8,27^. Due to its ease of use, the minimal computational power required, and because such methods can yield highly accurate and consistent segmentation results, there will be instances where global or local threshold methods are the superior method of choice. For example, even in this study, there was no significant differences in segmentation performance between the Yen global threshold and the vertebral body U-Net on the vertebral body dataset which demonstrates an instance where a simple global threshold may generate an accurate segmentation. However, threshold methods are inherently limited, as such approaches only consider greyscale variations in the surrounding voxels to identify bone voxles^34^. The advantage of a supervised deep learning approach is that the model weights are optimized and effectively “learn” from the input data manually created by humans^35^. Visually, humans can take in far more complicated information, understanding the underlying bone architecture and perceiving patterns and greyscale variations to identify bone. Deep learning models use computational algorithms that are fine-tuned to recognize underlying nuances to segment the structure. The disadvantage of the FCNNs presented here is that while the basic architecture is known, neural networks are notoriously considered “black-box” approaches as it is difficult to understand the complex algorithms that make decisions ^36,37^. Fortunately, this an active area of research, and it is likely that methods to help scientific researchers understand the method of choice will exist in the near future^36^.

A secondary finding of this study was the wide range in Dice indices obtained from different slices but the same dataset using local and global Otsu threshold approaches. The wide range of Dice indices demonstrates that accuracy assessment of a few cross-sectional slices is likely not appropriate, on its own, to validate the segmentation method used for a study. In addition, both the global and local Otsu threshold had low Dice indices (global Otsu 0.356 – 0.445; local Otsu: 0.283 – 0.444). Previous work showed that Dice indices of 0.81 yield large percent differences in bone microstructure measures relative to ground truth images (i.e., cortical BVF 24%, trabecular BVF 39%, trabecular thickness 21.7%)^38^. Future work will focus on identifying the critical DICE value needed to insure meaningful and accurate microstructural measures from the data. However, regardless of the segmentation method used (e.g., global threshold, local threshold, atlas-based, deep learning), the authors suggest that the method should be quantified and compared to a ground-truth test set. Considering that the segmentation method can have a significant effect in quantifying trabecular architecture, the standard practice of reporting and quantifying bone microstructure needs reconsideration to better ensure reproducibility in future studies.

The novelty of this study was the success of segmenting bone from multiple bone scans. Despite some similarities (e.g., both high resolution and cadaveric specimens), the vertebral body and femoral neck scans differed in the anatomical region, acquisition parameters, and pipelines used for pre-processing. The extent of variation in bones, scanning systems, and acquisition parameters that can be trained on a network and retain high performance is not yet known. However, our results indicate that there is indeed great potential for training an expanded CNN on disparate data. Future work will evaluate whether the trained network can perform well on bone scans from different specimens (e.g., rats and mice) more frequently seen in musculoskeletal research. An interesting finding of this study was that additional training with unlike data did not significantly change the segmentation quality on either the femoral neck or vertebral body dataset compared to the original FCNNs. However, the combined U-net had a higher Dice index for femoral neck segmentation (0.963) that for the vertebral body (0.912) dataset. Despite balanced classes in terms of training and testing of the two datasets, the learned features appear to be stochastically skewed to perform better on the femoral neck dataset. The reason for this outcome is not understood. However, the femoral neck data was acquired at an increased resolution, and thus had better delineation of bone-to-background boundaries. Perhaps the femoral neck data is simply a better fit for CNN’s architecture given its original design. Regardless, network training is an iterative process minimizing the loss function and does not consider the type of imaging data considered. Future work will develop models optimized to perform equally well on all datasets considered via the use of custom loss functions, altered hyperparameters, and data augmentation.

Despite success with training FCNNs for segmentation, some limitations need to be addressed. Time and effort were spent on manually creating the GT dataset for the bone/background FCNNs (3 hours/slice). However, efficiency may have been improved if a smaller GT dataset was used and augmentation was added to more effectively represent the scan variability (e.g., adding noise and shifting the histogram). A second limitation of this study was that this network was a trained 2-D FCNN which was applied on a slice-by-slice basis to segment a 3-D structure. Theoretically, there are several advantages to using a network algorithm capable of taking input from all dimensions to determine a voxel type (i.e. accurately assessing partial volume effects). Multiple studies have demonstrated that a 3-D FCNN outperforms a 2-D FCNN in terms of segmentation accuracy^39,40^. However, given the nature and size of the scans in this study, it was not feasible to train a network with 3-D dimensional GT data. Third, the hyper-parameters were selected via manual selection with systematic testing. Instead, several automated methods exist (e.g., grid search, random search, and Bayesian optimization) and could be applied to concisely and accurately determine the hyper-parameters that optimize model performance^41^. Also, other tensor-based approaches for segmentation provide promise, but the feasibility of CT scans has yet to be assessed^42,43^. Finally, our FCNNs were trained to segment entire cross sections, even though microstructural analyses are typically quantified on bone biopsies due to the field-of-view to resolution limitations with micro-CT^1^. A comparison of segmentation from these volumes may better represent segmentation differences in the threshold methods and FCNN performance ^8^. However, prior work demonstrated that a FCNN trained on entire cross-sections of bone performs similarly as the Otsu on bone cubes extracted from both the femoral neck and the femoral head^38^. Despite limitations, three FCNNs have been created that can accurately segment bone from a background in the vertebral body and the femoral neck.

In conclusion, three FCNNs were trained to accurately segment bone from background in nano-CT and micro-CT scans of the femoral neck and vertebral body that have both biological and scanning system variability. This current study was limited in scope as the data used was from only two resolutions, two anatomical locations, and two imaging technologies. Extending the network’s capability to include quality segmentation of clinical scans and more anatomical sites would increase its useful capabilities. FCNNs developed in this study are available for use on Dragonfly 4.0 and provide a reproducible, completely automated, approach for segmentation. Our FCNNs accurately quantify architectural results and provide a novel solution to improve outcome rigor and reproducibility by minimizing thresholding errors in musculoskeletal research.

## Acknowledgements

Research reported in this publication was supported by the National Institute of Arthritis and Musculoskeletal and Skin Diseases of the National Institutes of Health (KJJ: AR065424, AR069620, AR068452; TLB: AR064244). The content is solely the responsibility of the authors and does not necessarily represent the official views of the National Institutes of Health.

## Supplemental Figures

**Supplemental 1:**
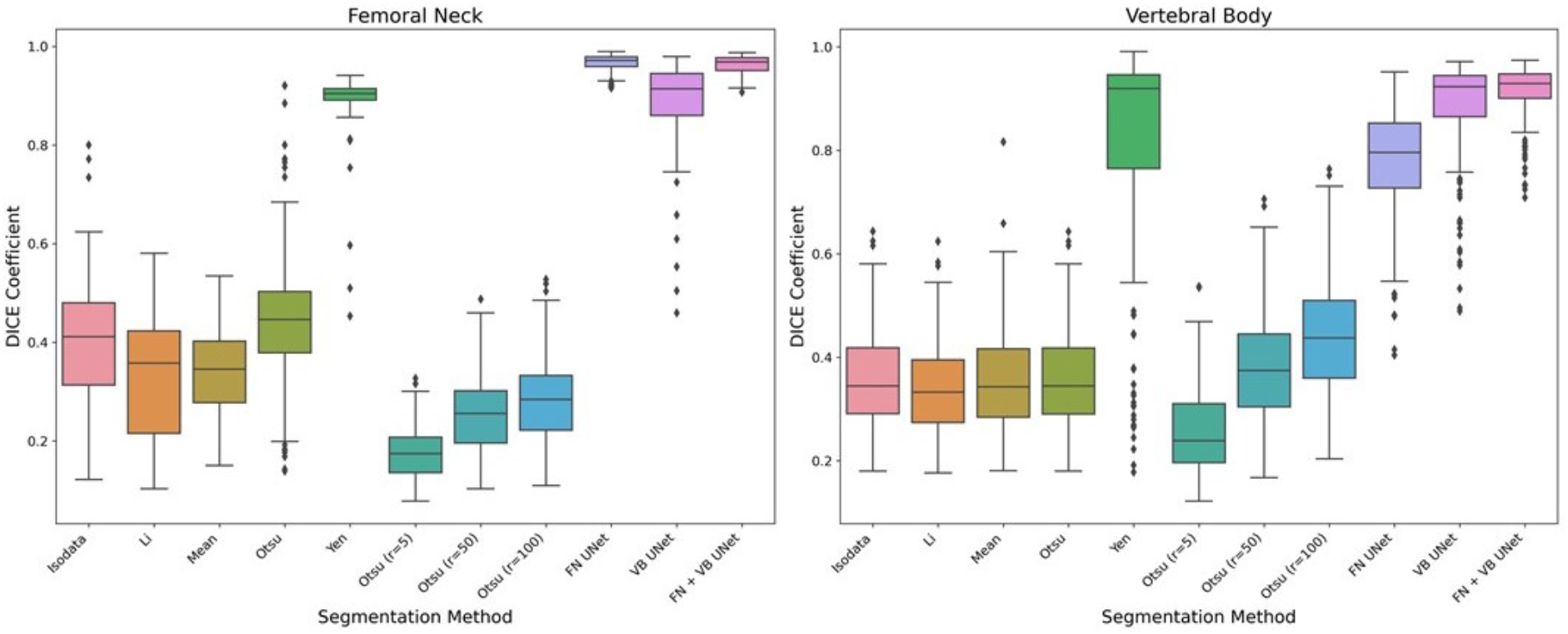
A boxplot of the DICE coefficients calculated on a slice-by slice basis for the global, local, and previously trained U-net segmentation methods relative to the ground truth for the femoral neck and vertebral body dataset.

